# PD-1-induced T cell exhaustion is controlled by a Drp1-dependent mechanism

**DOI:** 10.1101/2020.07.14.200592

**Authors:** Luca Simula, Valeria Cancila, Ylenia Antonucci, Alessandra Colamatteo, Claudio Procaccini, Giuseppe Matarese, Claudio Tripodo, Silvia Campello

**Author notes:** Department of Infection, Immunity, Inflammation (3I), Institut Cochin, INSERM U1016, CNRS UMR8104, F-75014, Paris, France.

## Abstract

PD-1 signalling downregulates the T cell response, promoting an exhausted state in tumor-infiltrating T cells, through mostly unveiled molecular mechanisms. Drp1-dependent mitochondrial fission plays a crucial role to sustain T cell motility, proliferation, survival and glycolytic engagement and, interestingly, such processes are exactly those inhibited by PD-1 in tumor-infiltrating T cells. Here we show that the signature of PD-1^pos^ CD8^+^ T cells infiltrating MC38-derived murine tumor mass is having downregulated Drp1 activity and more fused mitochondria, compared to PD-1^neg^ counterparts. Also, PD-1^pos^ lymphocytic elements infiltrating human colon cancer rarely express active Drp1. Mechanistically, PD-1 signalling directly prevents mitochondria fragmentation following T cell stimulation by downregulating Drp1 phosphorylation on Ser616, via regulation of the ERK1/2 and mTOR pathways. In addition, downregulation of Drp1 activity in tumor-infiltrating PD-1^pos^ CD8+ T cells seems to be a mechanism exploited by PD-1 signalling to reduce motility and proliferation of these cells. Overall, our data indicate that the modulation of Drp1 activity in tumor-infiltrating T cells may become a valuable target to ameliorate the anti-cancer immune response in future immunotherapy approaches.

## 1. INTRODUCTION

Programmed Cell Death-1 (PD-1) is a T cell surface receptor that down-regulates T cell activation and the immune response (Sharpe and Pauken, 2018). PD-1 signalling is activated by PD-1 interaction with its ligands PD-L1 and PD-L2, expressed on adjacent cells (Okazaki and Honjo, 2007), and it dampens signals originating from T Cell Receptor (TCR) and CD28, such as the activation of mammalian-Target-of-Rapamycin (mTOR) and Mitogen-Activated-Protein-Kinases (MAPK) pathways (Parry et al., 2005; Patsoukis *et al*., 2012). Besides being frequently observed in T cells during chronic infections (Jubel *et al*., 2020), activation of PD-1 signalling has also been widely reported in tumor-infiltrating T cells, contributing to their functional exhaustion and poor anti-tumor response (Wherry and Kurachi, 2015). Consistently, agonistic PD-1 antibodies efficiently reinvigorate tumor-infiltrating T cells, thereby ameliorating anti-tumor response (Iwai *et al*., 2017; Sun *et al*., 2018).

Mitochondria are central modulators of cellular bioenergetics, and their dynamic morphology is tightly linked to cell functions. Among the mitochondria-shaping proteins, Dynamin-Related Protein-1 (Drp1) is the main pro-fission protein and it is recruited from the cytosol to mitochondria thanks to post-translational modifications and to several receptors, such as Mff and Fis1 (Losón *et al*., 2013; Otera *et al*., 2013). Interestingly, mitochondria morphology is tightly linked to an optimal T cell functionality. Particularly, Drp1-dependent mitochondria fragmentation sustains T cell motility and proliferation, and effector T (T_eff_) cell apoptosis following TCR engagement (Simula *et al*., 2018, 2020). Also, Drp1-dependent mitochondria relocation at the immunological synapse controls the influx of calcium upon T cell activation (Baixauli *et al*., 2011), sustaining the cMyc-dependent upregulation of glycolytic enzymes (Simula *et al*., 2018; Wang *et al*., 2011), thus allowing the metabolic reprogramming required to cope with the increased bioenergetic demand of an activated T cell (Ma *et al*., 2019; Munford and Dimeloe, 2019). All these processes contribute to an optimal anti-tumor T cell response, which is indeed defective in T cells lacking Drp1 (Simula *et al*., 2018).

Of note, most of these Drp1-dependent processes are also down-regulated by PD-1 co-inhibitory signalling, especially when considering tumor-infiltrating T cells. Indeed, PD-1 signalling reduces T cell proliferation and motility (both of them requiring Drp1) (Patsoukis *et al*., 2012; Simula *et al*., 2018; Zinselmeyer *et al*., 2013), and it also promotes a shift from a glycolysis-based metabolism (supported by Drp1) toward an OXPHOS-based metabolism (requiring mitochondrial fusion) (Patsoukis *et al*., 2015). Given this striking inverse correlation, we asked whether PD-1 signalling may modulate Drp1 activity, and to what extent this modulation may down-regulate several processes in T cells. This point is of extreme importance, since the molecular mechanisms by which PD-1 regulates the aforementioned processes in T cells are not yet completely understood.

We here show that tumor-derived PD-1^pos^ CD8^+^ T cells exhibit a significant down-regulation of Drp1 activity and a more fused mitochondrial network. Mechanistically, PD-1 signalling prevents Drp1 activation following T cell stimulation by regulating its phosphorylation on Ser616 through the modulation of Extracellular-Regulated Kinase 1/2 (ERK1/2) and mTOR proteins. Also, we provide evidence that Drp1 down-regulation contributes to the reduced proliferation and motility of PD-1^pos^ tumor-infiltrating T cells and, as a consequence, we identify Drp1 as a possible target for future therapeutic approaches aiming at restoring anti-tumor response in PD1^pos^ exhausted CD8^+^ T cells.

## 2. RESULTS

### 2.1 PD1^pos^ CD8+ T cells from MC38-derived murine tumors show a reduced mitochondrial fission

To investigate if PD-1 signalling modulates the morphology of the mitochondrial network in tumor infiltrating T lymphocytes (TILs), we looked at PD-1^neg^ and PD-1^pos^ CD8^+^ T cells infiltrating an 18 days-old solid tumor mass derived from s.c. inoculation of MC38 cells (murine adenocarcinoma) in c57BL/6 WT mice. We took advantage of this tumor model, since being characterized by a high level of T cell infiltration (Simula *et al*., 2018). By comparing the expression levels of different mitochondria-shaping proteins in CD8+ TILs, we observed that PD-1^pos^ CD8^+^ TILs do not show altered levels of total Drp1 compared to PD-1^neg^ counterparts (Fig. 1a). However, we observed a specific down-regulation of Drp1 phosphorylation on its activating residue Ser616 (Fig. 1a), while the inhibitory phosphorylation on Ser637 did not vary. Also, we did not observe any differences in the expression of other main pro-fusion (Mfn1, Mfn2, Opa1) or pro-fission (Fis1, Mff) proteins (Fig. 1a).

**Figure 1.**
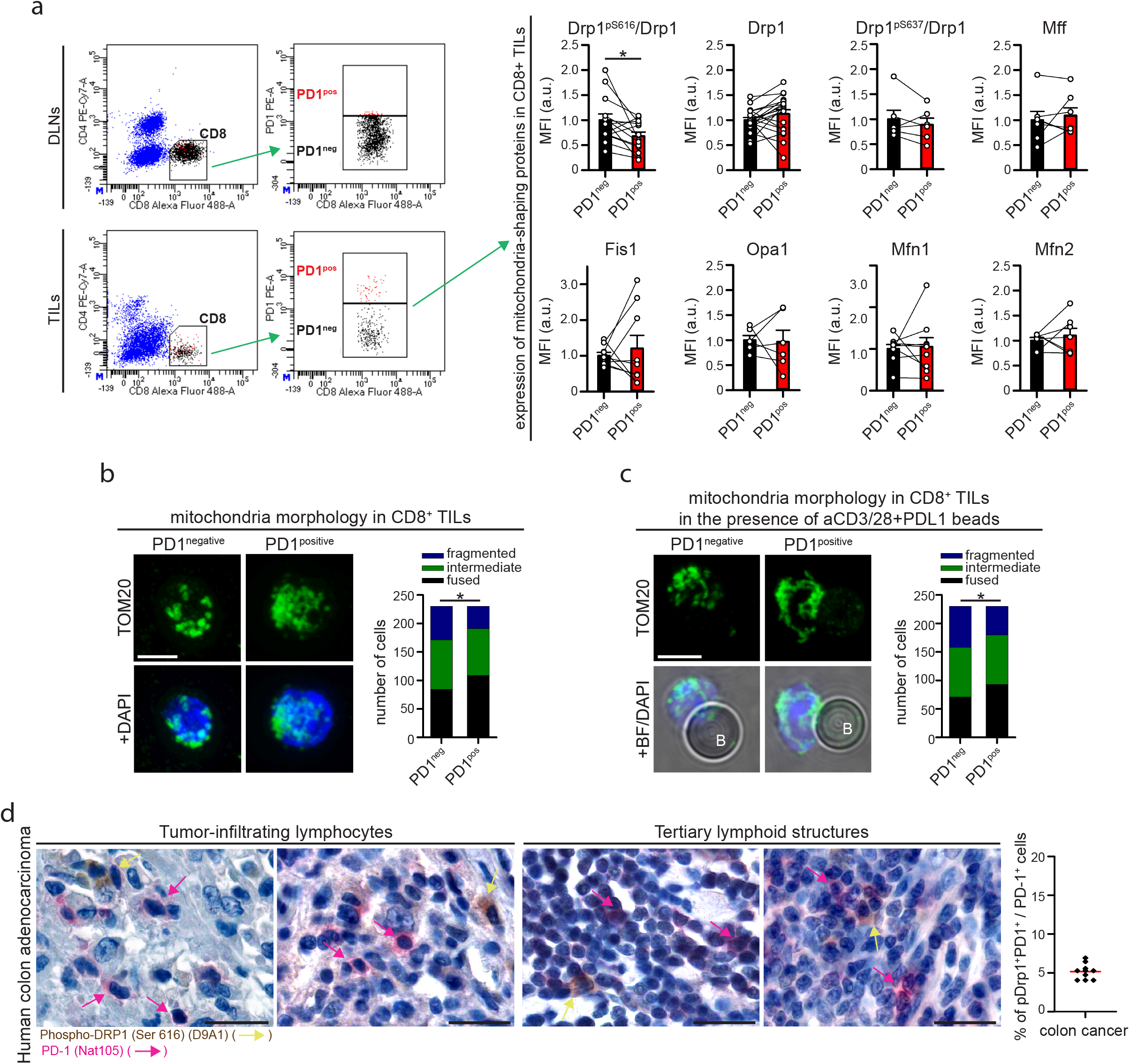
PD1^positive^ and PD1^negative^ CD8+ T cells within MC38-derived tumor microenvironment show different mitochondria morphologies. (**a**) Tumor-infiltrating lymphocytes have been isolated from 18 days-old MC38-derived tumor mass grown in WT c57BL/6 mice and the expression of the indicated mitochondria-shaping proteins have been evaluated into PD1^negative^ (PD1^neg^) and PD1^positive^ (PD1^pos^) CD8+ T cell subpopulations. Representative gating strategy to distinguish PD1^neg^ and PD1^pos^ CD8+ T cells is shown on the left. Graphs on the right indicate the normalized median fluorescence intensity (MFI) of the indicated proteins evaluated by intracellular flow cytometry in PD1^neg^ and PD1^pos^ CD8+ T cells from the same mice (pS616-Drp1 n=14; Drp1 n=20; Mfn1 and Fis1 n=9; Mfn2 and Mff n=7; Opa1 and pS637-Drp1 n=6; paired t-tests). (**b**-**c**) PD1^negative^ (PD1^neg^) and PD1^positive^ (PD1^pos^) CD44+ CD45+ CD8+ T cells have been sorted and purified form 18 days-old MC38-derived tumor mass grown in WT c57BL/6 mice. Gating strategy is shown in Supplemental Figure 1a. Mitochondria morphology was evaluated by immunofluorescence (anti-TOM20 staining) and upon z-stack reconstruction. In (b) cells have been fixed immediately after purification and processed for immunostaining. In (c) cells have been stimulated for 2h in the presence of beads coated with aCD3/28 Abs plus PDL1 and then fixed and processed for immunostaining. For each panel, representative images of the observed mitochondria morphologies are shown on the left, while graphs on the right show the distribution of cells into the indicated category according to mitochondria morphology in PD1n^eg^ and PD1^pos^ CD8+ T cells (n=230 cells each condition from 8 (b, unstimulated) or 10 (c, stimulated with beads) pooled mice; chi-square tests). (**d**) Representative microphotographs of double-marker immunohistochemistry for PD1 (rose; rose arrows) and Drp1-pSer616 (brown; yellow arrows) expression in lymphoid elements infiltrating human colon cancer. The graphs on the right indicates the percentage of double-positive pDp1^pos^PD-1^pos^ cells among all PD-1pos cells analysed (n=2731 PD-1^pos^ cells analysed from n=10 patients). Data are shown as mean ± SEM. Scale bar: 5µm in **b** and **c** and 50µm in **d**. Significance is indicated as follows: *=p<0.05.

Consistent with their reduced level of active Drp1, PD-1^pos^ CD8^+^ TILs show an altered morphology of mitochondria. Indeed, while PD-1^neg^ CD44^+^ CD8^+^ TILs show a fragmented network (as expected from activated T cells), mitochondria in PD-1^pos^ CD44^+^ CD8^+^ TILs are more fusion-prone (Fig. 1b, Supplementary Fig. 1a). Interestingly, this was observed also in sorted CD8^+^ TILs stimulated for 2h with beads coated with anti-CD3/28 antibodies plus PD-L1 (PD-1 ligand) before fixation (Fig. 1c), suggesting that such an altered mitochondria morphology may also influence T cell activation upon antigen encounter in a PD-L1-rich microenvironment (such as a tumor mass).

Last, we extended these observations to a homologue human tumor context, by staining moderately differentiated (G2) human colon carcinoma sections with anti-PD-1 and anti-Drp1-pSer616 antibodies. Of note, we found that tumor-infiltrating lymphocyte elements almost never co-express PD-1 and active Drp1. Indeed, only ca. 5% of PD-1^pos^ elements were also positive for Drp1^pSer616^ (Fig. 1d). Also, by staining sections with anti-CD8 antibody, we confirmed that almost all these PD-1^pos^ elements are CD8^+^ T cells (Supplementary Figure 1b). This suggests that also in a human tumor context PD-1^pos^ CD8^+^ T cells have downregulated Drp1 activity.

In sum, tumor-infiltrating PD-1^pos^ CD8^+^ T cells show a tendency toward a more interconnected mitochondria morphology, associated with a reduced activation of Drp1.

### 2.2 – PD-1 signalling prevents Drp1 activation and mitochondria fragmentation in both murine and human T cells, upon *in vitro* stimulation

To investigate whether such an altered Drp1 expression in PD-1^pos^ CD8^+^ TILs is directly caused by PD-1 activation, we switched to an *in vitro* system to specifically modulate PD-1 signalling. To this aim, we stimulated *in vitro* T cells isolated from spleen of WT mice with beads coated with anti-CD3/28 plus BSA (thereafter aCD3/28-beads) or anti-CD3/28 antibodies plus PD-L1 (thereafter aCD3/28-PDL1-beads) for 48h. As expected, concomitant activation of PD-L1/PD-1 axis during T cell stimulation dampens activation of both mTOR and ERK1/2 (Fig. 2a) (Parry *et al*., 2005; Patsoukis *et al*., 2012). Interestingly, we also observed that Drp1 phosphorylation on Ser616 is strongly reduced by the engagement of PD-1 signalling (Fig. 2a), while no significant difference was observed for other mitochondria-shaping proteins (Fig. 2b). In line with this, while T cells engage the fragmentation of mitochondria upon activation (Buck *et al*., 2016; Simula *et al*., 2018, 2020), in the presence of PD-L1 the same cell type retains a mitochondria morphology more similar to unstimulated counterparts (Fig. 2c). Of note, mitochondria of murine activated T cells might characteristically appear slightly swollen when fragmented (Fig. 2c), still being fully functional (Supplementary Fig. 2a, b).

**Figure 2.**
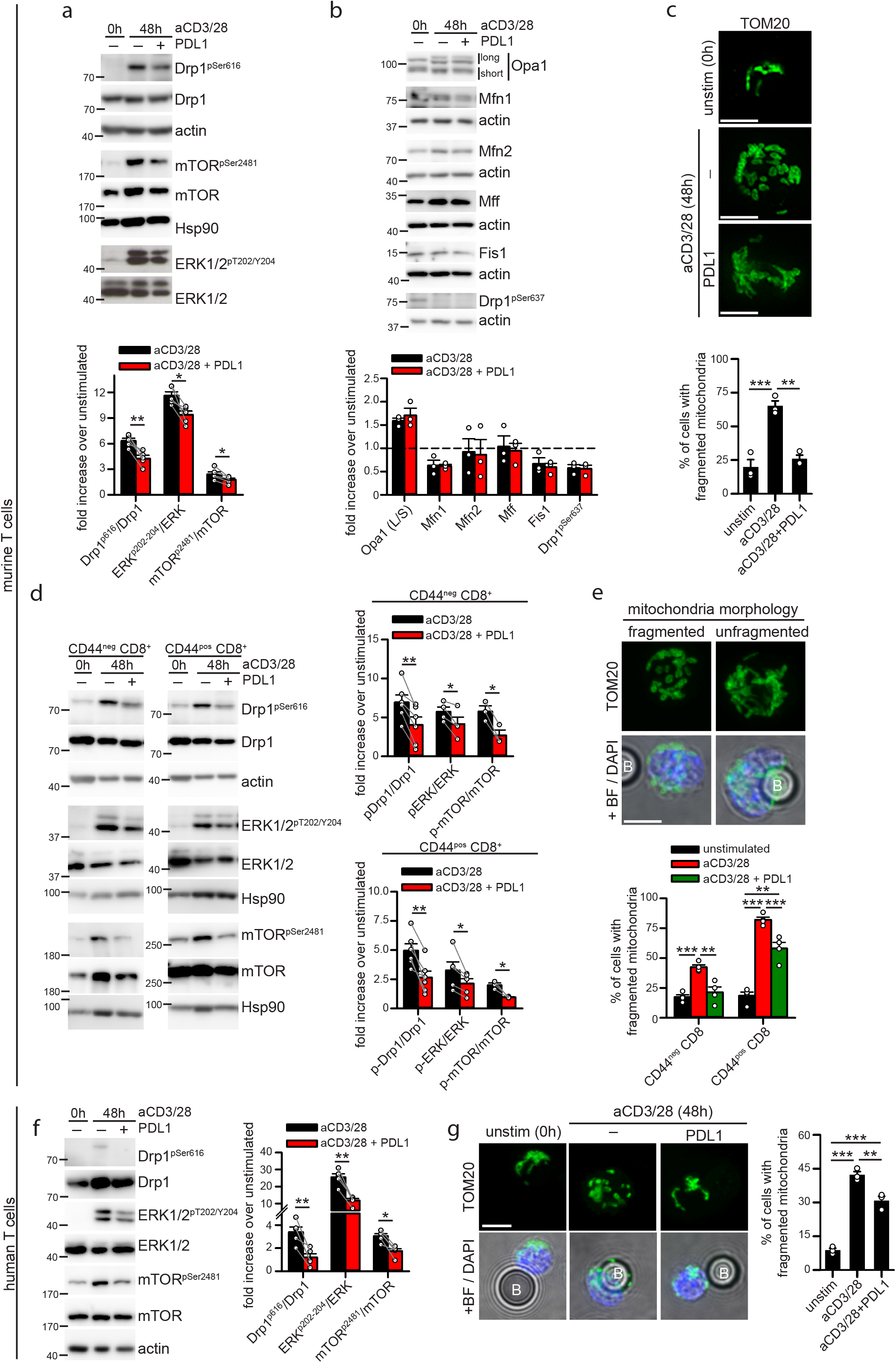
PD1 signaling downregulates Drp1-dependent mitochondria fragmentation in murine and human T cells. (**a**-**b**) Murine T cells have been isolated from spleen of WT c57BL/6 mice and left unstimulated (0h) or stimulated for 48h with anti-CD3/28- or anti-CD3/28-PDL1-beads. The expression level of the indicated (phospho)-proteins has been evaluated by western blot (a n=5, paired t-tests; b n=3). (**c**) Representative immunofluorescence images showing the mitochondrial network (anti-TOM20 staining) in murine T cells isolated and stimulated as in (a). Quantification of the percentage of cells showing fragmented mitochondria in each condition is reported in the graph below (n=3). (d) Murine CD44^neg^ (naïve) and CD44^pos^ (antigen experienced) CD8+ T cells have been isolated from spleen of WT c57BL/6 mice and left unstimulated (0h) or stimulated for 48h with anti-CD3/28- or anti-CD3/28-PDL1-beads. The expression level of the indicated (phospho)-proteins has been evaluated by western blot in both CD8+ T cell subsets (pDrp1 n=6; pERK naive n=4; pERK memory n=5; p-mTOR n=3; paired t-tests). (**e**) Immunofluorescence images showing representative mitochondrial morphologies (anti-TOM20 staining; “B” indicates a bead) observed in CD44^neg^ and CD44^pos^ CD8+ T cells isolated and stimulated as in (e). Quantification of the percentage of cells showing fragmented mitochondria in each condition is reported in the graph below (n=4). (**f**) Human T cells have been isolated from peripheral blood and left unstimulated (0h) or stimulated for 48h with anti-CD3/28- or anti-CD3/28-PDL1-beads. The expression level of the indicated (phospho)-proteins has been evaluated by western blot (n=5; paired t-tests). (**g**) Representative immunofluorescence images showing the mitochondrial network (anti-TOM20 staining; “B” indicates a bead) in human T cells isolated and stimulated as in (f). Quantification of the percentage of cells showing fragmented mitochondria in each condition is reported in the graph on the right (n=3). Data are shown as mean ± SEM. Scale bar: 5µm in **c, e** and **g**. Significance is indicated as follows: *=p<0.05; **=p<0.01; ***=p<0.001.

In addition, such an absence of fragmented mitochondria in PD-L1-stimulated T cells is not due to ongoing mitophagy (which could hypothetically promote the clearance of the small and fragmented mitochondria) (Supplementary Fig. 2c).

Next, we asked whether such PD-1-dependent regulation of Drp1 is restricted or not to some specific T cell subpopulation. Therefore, we looked more closely at naïve (CD44^neg^) and antigen experienced (CD44^pos^) CD8^+^ T cell subpopulations, which, once activated, may undergo exhaustion within the tumor microenvironment. Of note, although naïve CD8+ T cells do not express PD-1 on their cell surface, they rapidly acquire PD-1 expression 12h after stimulation with aCD3/28-beads (as also observed for naïve CD4^+^ T cells), and the concomitant presence of PD-L1 ligand does not affect such upregulation (Supplementary Fig. 2d). We thus isolated both subpopulations from the spleen of WT mice and stimulated them *in vitro* with aCD3/28- or aCD3/28-PDL1-beads for 48h, which is the optimal time point to detect Drp1 phosphorylation, especially in naïve CD8^+^ T cells (Supplementary Fig. 2e). Interestingly, we found that engagement of PD-1 signalling during cell activation dampens Drp1 phosphorylation and mitochondria fragmentation in both CD44^pos^ and CD44^neg^ CD8^+^ subsets (Fig. 2d, e), in parallel with a reduced activation of ERK and mTO R pathways (Fig. 2d).

Last, a PD-1-dependent Drp1 modulation was also observed in human T cells isolated from healthy donors’ peripheral blood (hPBT). Similar to their murine counterpart, hPBT cells stimulated for 48h with aCD3/28-PDL1-beads show Drp1 phosphorylation on Ser616 (Fig. 2f) and fragmented mitochondria (Figure 2g) significantly lower than aCD3/28-stimulated hPBTs, which also display higher mTOR and ERK1/2 activation (Fig. 2f).

These data indicate that co-stimulation of PD-1 signalling during T cell activation prevents Drp1 phosphorylation and mitochondria fragmentation both in mice and humans.

### 2.3 – PD-1 signalling downregulates Drp1 activation by modulating mTOR and ERK pathways

We aimed at better investigating the molecular pathways linking PD-1 activation to the downregulation of Drp1 activity. It is well established that PD-1 dampens the signalling pathways originating from TCR and CD28 activation (Parry *et al*., 2005; Patsoukis *et al*., 2012). Recently, an *in vitro* protocol based upon subsequent cycles of CD3/CD28 stimulation (see Methods for details) has been developed to induce a TCR/CD28-hyporesponsive state in CD8^+^ T cells, thus mimicking an exhausted-like condition of diminished proliferative and cytotoxic potential, without the need of a concurrent engagement of inhibitory co-receptors, such as PD-1 (Dunsford *et al*., 2020). Therefore, we took advantage of this protocol to understand whether PD-1 may regulate Drp1 phosphorylation mainly through its inhibition of TCR/CD28-derived signalling or by other mechanisms. To this aim, we compared Drp1 phosphorylation and mitochondria morphology in TCR/CD28 responder T_eff_-like cells (i.e. stimulated up to two times with aCD3/CD28) and non-responder T_ex_-like cells (i.e. the same cells stimulated three or more times). Interestingly, we observed that Drp1 phosphorylation and fragmentation of mitochondria are strongly reduced after three or more cycles of aCD3/CD28 stimulation compared to healthy effector T cells stimulated up to two times (Fig. 3a, b). Therefore, a reduced strength of TCR/CD28-derived signalling is sufficient to account for the effects observed on Drp1 and mitochondria morphology in an exhausted-like T cell. These data also suggest that modulation of the TCR/CD28 signalling may represent the main mechanism by which PD-1 could regulate the mitochondrial morphology in T cells.

**Figure 3.**
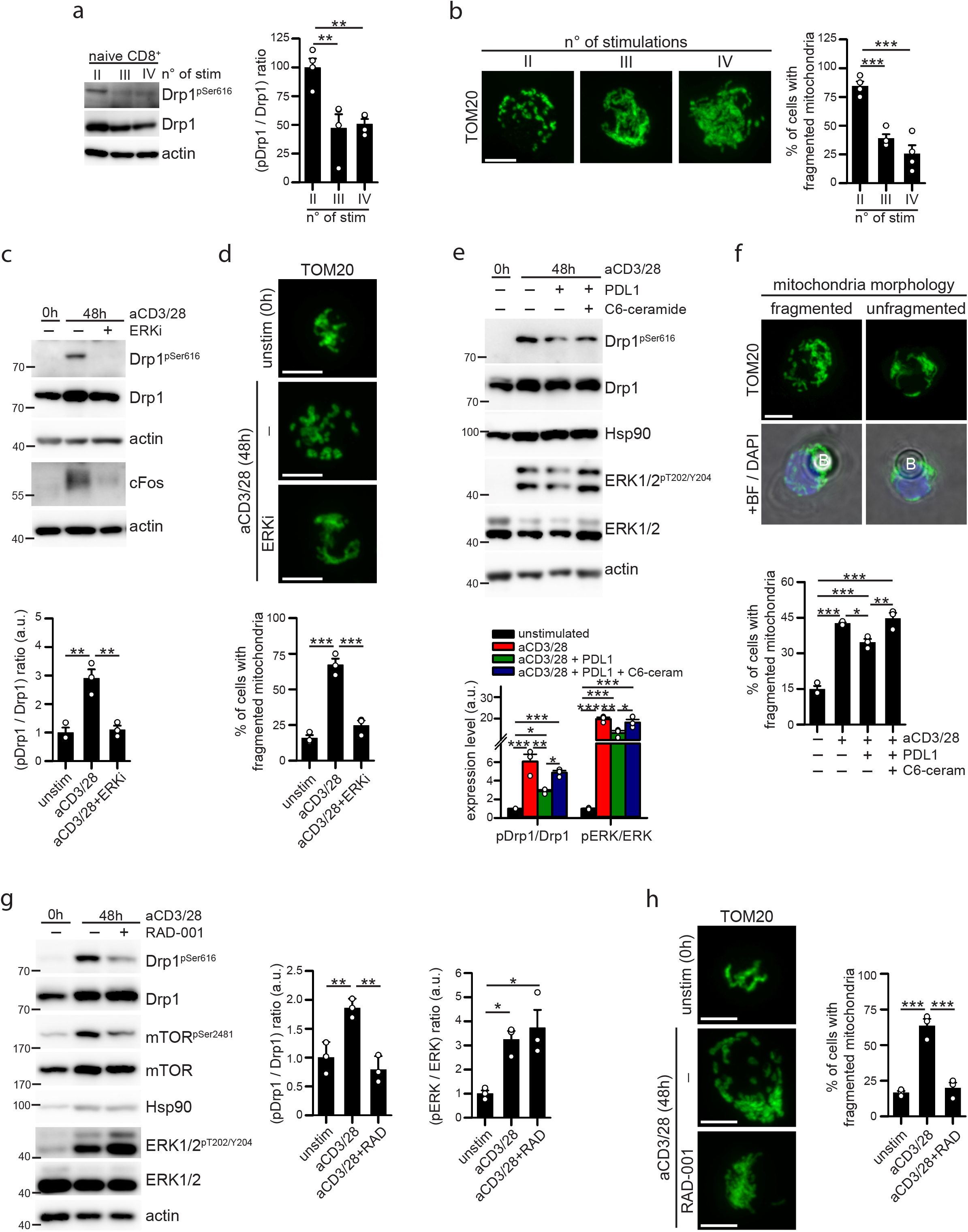
PD1 signaling down-modulates Drp1 activity via ERK and mTOR pathways. (**a**) Murine CD44^neg^ naïve CD8+ T cells have been isolated from spleen of WT c57BL/6 mice and stimulated up to 4 times with plate-coated anti-CD3 Ab plus soluble anti-CD28 for 24h. After each stimulation, cells were left to recover 6 days in IL2-containing medium before the next stimulation. The expression level of the indicated (phospho)-proteins has been evaluated by western blot immediately after the second (II), third (III) and fourth (IV) stimulation (n=4). (**b**) Representative immunofluorescence images showing the mitochondrial network (anti-TOM20 staining) in murine CD44^neg^ naïve CD8+ T cells stimulated as in (c). Quantification of the percentage of cells showing fragmented mitochondria in each condition is reported in the graph on the right (n=4). (**c**-**d**) Murine T cells have been isolated from spleen of WT c57BL/6 mice and stimulated for 48h with anti-CD3/28-coated beads in presence or not of 30µM FR180204 (ERK inhibitor: ERKi). In (c) is reported the expression level of the indicated (phospho)-proteins evaluated by western blot and quantified in the graph below (n=3). In (d) are reported representative immunofluorescence images showing the mitochondrial network (anti-TOM20 staining) in murine T cells. Quantification of the percentage of cells showing fragmented mitochondria in each condition is reported in the graph below (n=3). (**e**-**f**) hPBT cells have been isolated from spleen of WT c57BL/6 mice and stimulated for 48h with anti-CD3/28- or anti-CD3/28-PDL1-beads in presence or not of 10µM C6-ceramide (ERK activator). In (e) is reported the expression level of the indicated (phospho)-proteins evaluated by western blot and quantified in the graph below (n=3). In (f) are reported representative immunofluorescence images showing the mitochondrial network (anti-TOM20 staining, “B” indicates a bead) in hPBT cells. Quantification of the percentage of cells showing fragmented mitochondria in each condition is reported in the graph below (n=3). (**g**-**h**) Murine T cells have been isolated from spleen of WT c57BL/6 mice and stimulated for 48h with anti-CD3/28-coated beads in presence or not of 10nM RAD-001 (mTOR inhibitor). In (g) is reported the expression level of the indicated (phospho)-proteins evaluated by western blot and quantified in the graphs on the right (n=3). In (h) are reported representative immunofluorescence images showing the mitochondrial network (anti-TOM20 staining) in murine T cells. Quantification of the percentage of cells showing fragmented mitochondria in each condition is reported in the graph on the right (n=3). Data are shown as mean ± SEM. Scale bar: 5µm in **b, d, f** and **h**. Significance is indicated as follows: *=p<0.05; **=p<0.01; ***=p<0.001.

To identify more specifically the TCR/CD28-dependent targets of PD-1 which could account for its modulation of Drp1 activity, we reasoned that PD-1 is known to dampen both PI3K/Akt/mTOR pathway downstream of CD28 activation (Parry *et al*., 2005) and MAPK/ERK pathway downstream of TCR engagement (Patsoukis *et al*., 2012), as indeed confirmed by our results (Fig. 2a, d, f). Since both these pathways have been reported to modulate Drp1-dependent mitochondria fragmentation (Kashatus *et al*., 2015; Morita *et al*., 2017), we tested whether their inhibition is sufficient to reduce Drp1 activation. First, ERK1/2 inhibitor FR180204 (ERKi) indeed prevents both Drp1 phosphorylation and mitochondrial fragmentation in activated T cells (Fig. 3c, d). Further, we rescued ERK1/2 activity down-stream of PD-L1/PD-1 engagement by using low doses of C6-ceramide (to avoid apoptosis induction), a known activator of the ERK pathway (Raines *et al*., 1993). Of note, ceramide, which slightly increase ERK phosphorylation, rescues Drp1 phosphorylation and mitochondria fragmentation in PD-1-engaged hPBT cells during activation (Fig. 3e, f). Next, also the mTOR inhibitor RAD-001 (Sedrani *et al*., 1998) prevents both Drp1 phosphorylation and mitochondria fragmentation during T cell activation (Fig. 3g, h). Interestingly, while ERK1/2 is known to directly phosphorylate Drp1 on Ser616 (Kashatus *et al*., 2015), it is currently unknown how mTOR may regulate Drp1 activity. An extensive crosstalk between mTOR and ERK pathways has been frequently reported (Darvishi *et al*., 2017; Mendoza *et al*., 2011). Therefore, mTOR may regulate Drp1 in T cells via the modulation of ERK1/2. However, we here observed that a low dose (10nM) of mTOR inhibitor is sufficient to prevent Drp1 phosphorylation without affecting ERK1/2 activity (Fig. 3g). Therefore, at least in T cells, mTOR seems to regulate Drp1 in a ERK1/2-independent way.

In sum, activation of PD-1 signalling during T cell stimulation reduces Drp1 phosphorylation and mitochondria fragmentation presumably through an inhibition of the ERK and mTOR pathways downstream of TCR/CD28 signalling.

### 2.4 Drp1 is required for an efficient reduction of tumor growth mediated by anti-PD-1 therapy

Given the importance of Drp1 in regulating multiple processes in T cells, we asked whether the ability of anti-PD-1 therapy to reduce solid tumor growth requires the restoration of Drp1 activity in tumor-infiltrating PD-1^pos^ CD8^+^ T cells. To answer this point, we analysed s.c. MC38-derived tumors in mice, whose growth *i)* is significantly reduced by treatment with anti-PD-1 Abs (Grasselly *et al*., 2018), and *ii)* requires a functional Drp1 in T cells (Simula *et al*., 2018). We s.c. inoculated MC38 cells into both control (Drp1^fl/fl^) and Drp1 conditional-KO (Drp1^fl/fl^ Lck:cre+, T cell-restricted Drp1 ablation, indicated as Drp1cKO) mice, which we characterized previously (Simula *et al*., 2018). After one week, we treated these mice with either anti-IgG (control) or anti-PD-1 antibody every 2/3 days for up to 10 days (Fig. 4a). Of note, we found that anti-PD-1 treatment is much more efficient in reducing tumor growth in control mice than in mice where Drp1 was absent in T cells (Drp1-cKO) (Fig. 4b), even when correcting for the larger tumor volume in Drp1-cKO mice compared to control mice (Simula *et al*., 2018). Even so, we observed a drop from 60% to ca. 20% in the efficacy of anti-PD-1 therapy in Drp1-cKO mice (Fig. 4c). In addition, we found that the strong reduction in tumor growth by anti-PD-1 treatment in control (Drp1^fl/fl^) mice correlates with a rescue of Drp1 phosphorylation in PD1^pos^ CD8^+^ T cells to a level comparable to PD1^neg^ CD8^+^ T cells (Fig. 4d). On the contrary, in Drp1-cKO mice (where Drp1 is absent and therefore cannot be “rescued”), anti-PD-1 treatment is much less efficient in reducing tumor growth. Overall, these data suggest that the restoration of Drp1 activity in PD1^pos^ CD8^+^ TILs is a key step for the effectiveness of the anti-PD-1 therapy.

**Figure 4.**
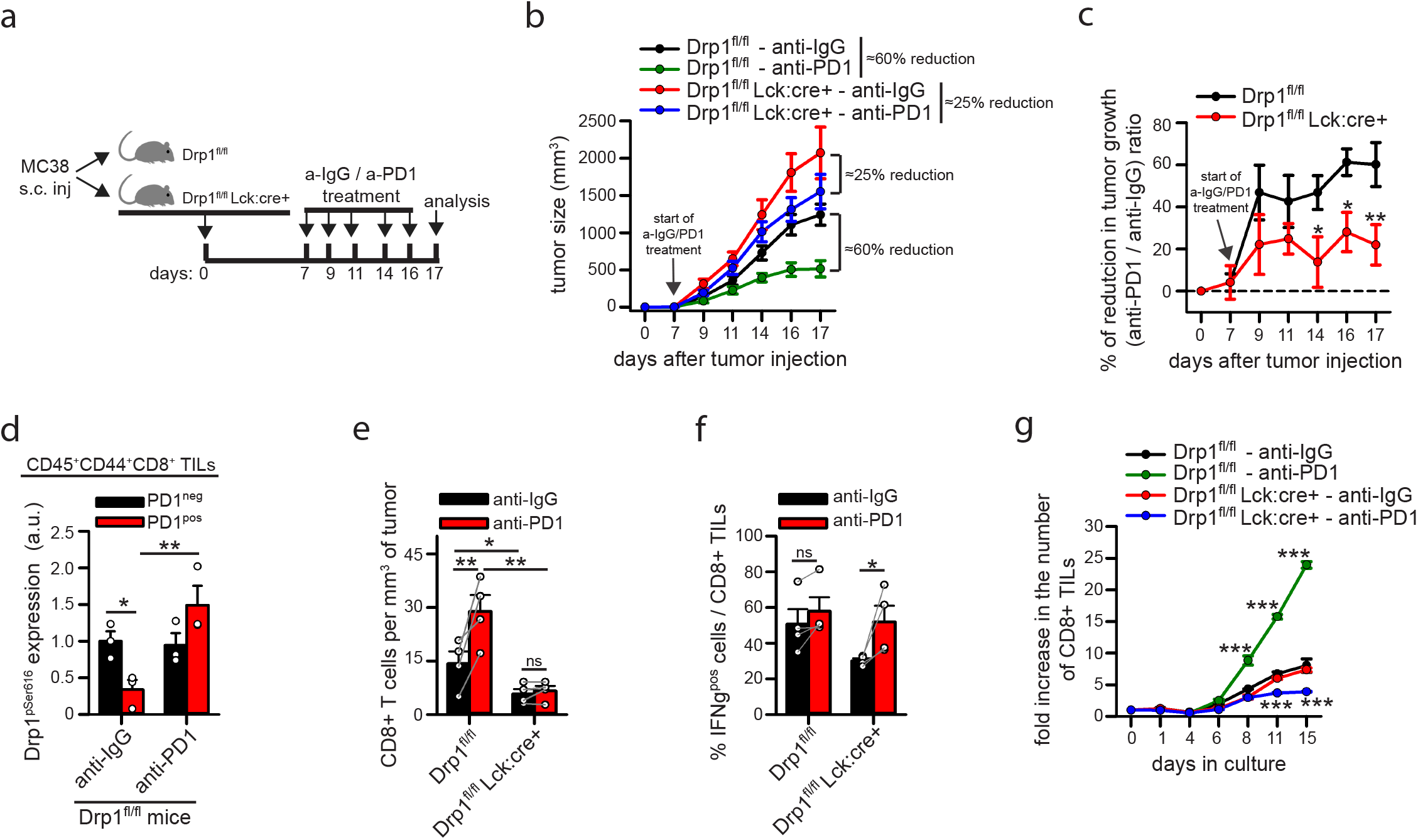
The downregulation of Drp1 activity in tumor-derived PD-1^pos^ T cells contributes to tumor growth. **(a)** Schematic representation of the experimental plan. (**b**) Size of MC38-derived tumors grown for the indicated days in control Drp1^fl/fl^ or conditional-KO Drp1^fl/fl^Lck:cre+ mice inoculated with anti-IgG or anti-PD1 antibodies as indicated in (a) (Drp1^fl/fl^ anti-IgG n=8; Drp1^fl/fl^ anti-PD1 and Drp1^fl/fl^Lck:cre+ anti-IgG n=12; Drp1^fl/fl^Lck:cre+ anti-PD1 n=14). (**c**) Relative percentage of the reduction in tumor growth (anti-PD1 / anti-IgG ratio) calculated from data in (b) for control Drp1^fl/fl^ or conditional-KO Drp1^fl/fl^Lck:cre+ mice (Drp1^fl/fl^ n=12; Drp1^fl/fl^Lck:cre+ n=14). (**d**) Quantification of the Drp1-pSer616 median fluorescence intensity (MFI) evaluated by intracellular flow cytometry in CD45+ CD44+ CD8+ PD1^neg^ and PD1^pos^ TILs from control Drp1^fl/fl^ mice. The MFI value obtained in corresponding TILs from conditional-KO Drp1^fl/fl^Lck:cre+ mice has been used as negative control and subtracted from the corresponding population in Drp1^fl/fl^ mice to obtain pSer616-Drp1 MFI data reported in the graph (n=3). (**e**) Absolute number of CD8+ T cells per mm^3^ of tumor collected from MC38-derived tumor masses grown for 17 days as described in (a) (n=4; Two-Way ANOVA on repeated measurements). (**f**) TILs have been isolated from MC38-derived tumor masses grown for 17 days as described in (a) and stimulated in vitro for 4h to evaluate IFNγ production. Percentage of IFNγ^pos^ cells among CD8+ T cells in each condition is reported in the graph (n=4; Two-Way ANOVA on repeated measurements). (**g**) TILs have been isolated from MC38-derived tumor masses grown for 17 days as described in (a) and cultured *in vitro* for the indicated days in presence of IL-2, IL-7 and IL-15 cytokines. Quantification of the fold increase in the absolute number of CD8+ T cells per day (starting value at day 0 = 1) is reported in the graph. Significance is indicated for each single point only if that day the population differs statistically from all the other 3 populations (n=3). Data are shown as mean ± SEM. Significance is indicated as follows: *=p<0.05; **=p<0.01; ***=p<0.001.

Next, we tried to get more insight into the cellular processes requiring the rescue of Drp1 activity during anti-PD-1 therapy. Of note, while anti-PD-1 treatment increases the number of CD8^+^ TILs recovered from the tumor mass (per mm^3^), this effect is not observed in Drp1-cKO mice (Fig. 4e), indicating that Drp1 is required to mediate such anti-PD-1-dependent increase in CD8^+^ TILs accumulation. On the contrary, PD-1-dependent regulation of IFNγ production does not involve Drp1 (Fig. 4f). We previously reported that Drp1 is also required to sustain T cell clonal expansion after stimulation, both *in vitro* and *in vivo* (Simula *et al*., 2018). Therefore, we asked whether the inability of Drp1-KO CD8^+^ TILs to increase their number within the tumor mass upon anti-PD-1 treatment may depend or not on their impaired proliferation. To this aim, we isolated TILs from MC38-derived tumor masses grown in control or Drp1-cKO mice (treated with anti-IgG or anti-PD-1) and let them expand *in vitro* in the presence of IL-2, IL-7 and IL-15 cytokines. Interestingly, while the anti-PD-1 treatment significantly increases fold expansion of control CD8^+^ T cells, this effect is completely lost in Drp1-KO CD8^+^ T cells (Fig. 4g). These data suggest that Drp1 inhibition may be at least one of the mechanisms by which PD-1 signalling reduces proliferation of PD1^pos^ CD8^+^ TILs.

In sum, a functional Drp1 is required for the efficacy of the anti-PD-1 treatment in reducing MC38-derived tumor growth in mice. Also, PD-1 signalling may mediate the reduction in CD8^+^ TILs proliferative potential via Drp1 downregulation.

### 2.5 Reduced motility of PD1_pos_ and *in vitro* exhausted-like T cells correlates with altered Drp1-dependent mitochondrial remodelling

Besides controlling T cell proliferation, Drp1 is required to sustain T cell motility by favoring mitochondria repositioning at the uropod, and it is directly phosphorylated on Ser616 in response to chemokine stimulation (Campello *et al*., 2006; Simula *et al*., 2018). Also, T cell motility is another process dampened by PD-1 signalling (Zinselmeyer *et al*., 2013). Therefore, in addition to controlling proliferation of PD1^pos^ CD8^+^ TILs, the PD-1-dependent downregulation of Drp1 activity may also account for their reduced motility, and its rescue being required to restore motility during anti-PD-1 therapy. To address this point, we isolated TILs from MC38-derived tumor masses grown in control or Drp1-cKO mice (treated with anti-IgG or anti-PD-1) and let them starve *in vitro*. Then, we assayed their migratory ability in response to serum by using the transwell migration assay. Interestingly, we found that anti-PD-1 treatment significantly increases motility of control PD1^pos^ CD8^+^ TILs, when compared to the motility of the same cells from anti-IgG-treated control mice (Fig. 5a). However, this effect is completely lost when looking at PD1^pos^ CD8^+^ TILs from Drp1-cKO mice, whose T cells lack Drp1 (Fig. 5a). Again, these data suggest that *i)* the rescue of Drp1 activity by anti-PD-1 treatment is required to restore TILs motility and *ii)* Drp1 inhibition may be one of the mechanisms by which PD-1 signalling reduces motility of PD1^pos^ CD8^+^ TILs.

**Figure 5.**
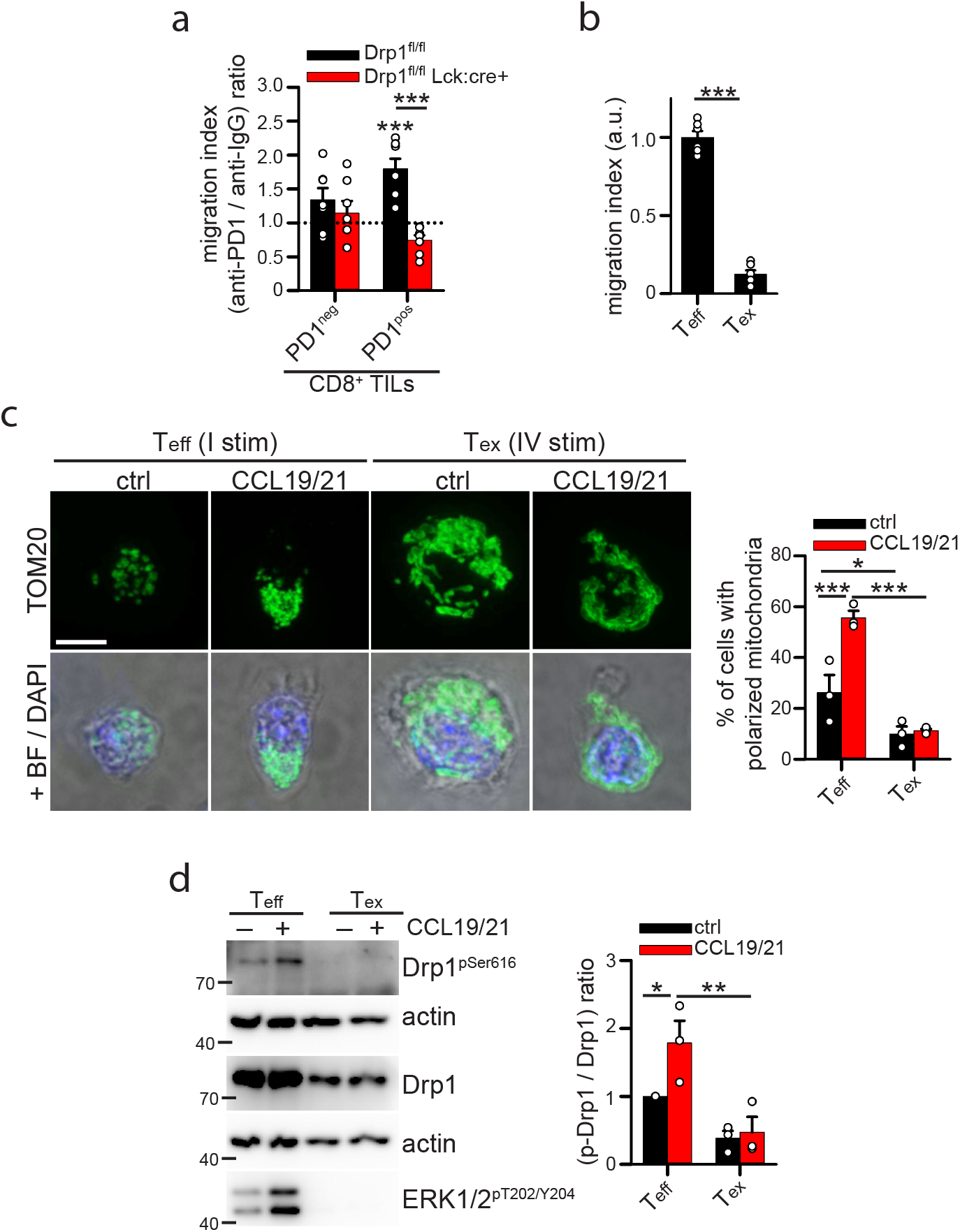
The downregulation of Drp1 activity contributes to the reduced motility of PD1^positive^ T cells. (**a**) TILs have been isolated from MC38-derived tumor masses grown for 17 days in control Drp1^fl/fl^ or conditional-KO Drp1^fl/fl^Lck:cre+ mice inoculated with anti-IgG or anti-PD1 antibodies as indicated in Figure 4a. Then, TILs were starved from serum for 2h and allowed to migrate in response to 10% Fetal Bovine Serum for 2h using transwell migration assay. The graph indicates the relative (anti-PD1 / anti-IgG) migration index calculated for PD1^neg^ and PD1^pos^ CD8+ TILs isolated from tumor-bearing control Drp1^fl/fl^ or conditional-KO Drp1^fl/fl^Lck:cre+ mice inoculated with anti-IgG or anti-PD1 antibodies as indicated in Figure 5a (n=7). (**b**-**d**) Murine exhausted T cells (Tex) have been generated in vitro through 4 cycles of a-CD3/28-mediated stimulation (24h) and IL2-mediated expansion (6 days) and compared to effector T cells (Teff) generated through a single cycle of stimulation and expansion. After the last 6 days in IL2-containing medium, cells have been starved from serum for 2h and then the following assays were performed. In (b) the migration index in response to CCL19/CCL21 gradient for 2h has been calculated using transwell migration assay (n=6). In (c) cells have been left to adhere for 30min to fibronectin-coated slides. Then, cells have been stimulated with 50nM CCL19 and 50nM CCL21 chemokines for 15min and then fixed and processed for immunostaining. Representative images showing the mitochondrial network (anti-TOM20 staining) in effector T (Teff) and exhausted T (Tex) cells are shown on the left. Quantification of the percentage of cells showing fragmented mitochondria in each condition is reported in the graph on the right (n=3). In (d) the cells have been stimulated with 50nM CCL19 and 50nM CCL21 chemokines for 15min in an eppendorf tube and then proteins were extracted. The expression level of the indicated (phospho)-proteins has been evaluated by western blot (n=3). Data are shown as mean ± SEM. Scale bar: 5µm in **c**. Significance is indicated as follows: *=p<0.05; **=p<0.01; ***=p<0.001.

To analyse in more details the modulation of Drp1 in functional and exhausted T cells during T cell migration, we switched to the aforementioned *in vitro* model to induce an exhaustion-like state in CD8^+^ T cells. As expected, exhausted-like (T_ex_) cells migrate much less than effector (T_eff_) CD8+ cells (Fig. 5b). Interestingly, we also observed that while functional T_eff_ cells correctly polarize the mitochondria at the uropod (Fig. 5c), a process requiring Drp1 (Simula *et al*., 2018), and show both Drp1 and ERK1/2 phosphorylation upon chemokine stimulation (Fig. 5d), in T_ex_ cells the mitochondrial network fails to rearrange (Fig. 5c), and Drp1 and ERK1/2 are unresponsive to chemokine stimulation (Fig. 5d).

Overall, these data indicate that PD-1 signalling may mediate the reduction in CD8^+^ TILs motility via Drp1 downregulation. Also, *in vitro*-induced exhausted-like T cells lack the ability to activate the Drp1-dependent mitochondria fragmentation upon chemokine stimulation, a key step normally required for T cell motility.

## 3. DISCUSSION

It has been reported that tumor-infiltrating T cell show an altered mitochondria functionality and morphology when compared with T cells from peripheral blood (Siska *et al*., 2017). However, the modulation of mitochondria morphology in different subpopulations of TILs has never been investigated before. Here, we report that in MC38-derived tumors PD1^pos^ tumor-infiltrating murine CD8+ T cells display a reduced activation of Drp1 and a more fused mitochondrial network when compared with PD1^neg^ counterparts. Of note, these data are shared also in a corresponding human context of colon tumor, in which tumor-infiltrating lymphocytic elements almost never co-express PD-1 and active Drp1. Mechanistically, we provided evidence that PD-1 signalling downregulates Drp1 activating phosphorylation on Ser616 (and consequently mitochondria fragmentation) via the inhibition of ERK1/2 and mTOR kinases.

Also, we explored the functional consequences of such PD-1-dependent downregulation of Drp1 activity in the tumor context. Of the highest importance, altogether, our data suggest that PD-1 signalling may exploit the downregulation of Drp1 activity to dampen some of the processes required for an optimal T cell functionality. In line with this, the restoration of Drp1 activity in TILs is strictly required for the effectiveness of the anti-PD-1 therapy. Specifically, Drp1 seems to play an important role in controlling both motility and proliferation of PD1^pos^ CD8+ TILs. Mechanistically, we observed that the Drp1-dependent mitochondria relocation at the cell rear-edge during cell migration, a phenomenon occurring in healthy effector CD8+ T cells (Simula *et al*., 2018), is completely lost in exhausted CD8+ T cells, which are unable to activate Drp1 upon chemokine stimulation. Our data are thus consistent with previous observations made in a persistent infection mouse model, in which the recovered motility of PD1^pos^ T cells upon anti-PD-1 treatment were associated with an increased activation of ERK (Zinselmeyer *et al*., 2013), a kinase known to regulate Drp1 in T cells (Simula *et al*., 2018, 2020). Therefore, downregulation of ERK/Drp1 axis may be exploited by PD-1 signalling to dampen T cell motility. Regarding the role of Drp1 in T cell proliferation, we previously reported that, in the absence of Drp1, T cells show an abnormal length of mitosis, due to the acquisition of aberrant centrosome morphologies (Simula *et al*., 2018), as observed also in cancer cells (Qian *et al*., 2012).

Moreover, we could speculate that the downregulation of Drp1 by PD-1 signalling may represent a mechanism exploited by exhausted TILs to modulate mitophagy and metabolism, too. First, a defective mitophagy in PD1^positive^ T cells has been reported to contribute to the exhausted phenotype of these cells (Yu *et al*., 2020). Interestingly, Drp1 may facilitate dismissal of small mitochondria via mitophagy (Twig *et al*., 2008). Therefore, the downregulation of Drp1 activity in PD1^pos^ CD8+ TILs may provide a mechanism to reduce the mitophagy rate in these cells, preventing the generation of small mitochondria that can be targeted to degradation. This is consistent with the observation of a higher mitochondrial mass in CD8+ TILs (Siska *et al*., 2017). Second, it is known that the engagement of PD-1 signalling downregulates glycolysis in T cells, at least partly by modulating mTOR pathway (Patsoukis *et al*., 2015). Since we previously reported that Drp1 interacts with mTOR to promote glycolysis upon T cell activation (Simula *et al*., 2018), our observation that PD1^pos^ CD8+ T cell downregulates Drp1 activation upon stimulation may provide a better understanding of the mechanistic basis for the downregulation of the glycolysis by PD-1 signalling.

Of note, Ogando *et al*. recently reported no variations in the rate of mitochondrial fragmentation in T cells unstimulated or stimulated in the presence or absence of PD-1 engagement (Ogando *et al*., 2019). However, fragmentation of the mitochondrial network upon T cell stimulation (independently of PD-1 signalling) has been reported extensively by us and others (Baixauli *et al*., 2011; Buck *et al*., 2016; Simula *et al*., 2018, 2020). We suppose that such discrepancy may rely partially on their use of the circularity as a parameter to estimate the rate of mitochondrial fragmentation. Indeed, we believe that such a parameter may not be suitable to compare mitochondria morphology in stimulated and unstimulated (naïve) T cells, since the former cells grow and enlarge significantly compared to the latter ones upon *in vitro* culture. Also, fused mitochondria in T cells appear more tangled due to the round shape of this non-adherent and small type of cells. In addition, Ogando *et al*. only analysed total levels of Drp1, without focusing on its specific phosphorylated residues (Ogando *et al*., 2019), which are more reliable indicators of Drp1 activation compared to the total protein amount.

In sum, our data indicate that downregulation of Drp1 mediated by PD-1 signalling may be required to attain an efficient inhibition of T cell response. Therefore, we dare to propose Drp1 as a therapeutic target to ameliorate exhausted T cell functionality during anti-cancer approaches, although drugs able to activate this protein need to be developed yet. Interestingly, CAR T cell-based approaches are currently being exploited for the treatment of solid cancers (Caruana *et al*., 2018), but they frequently fail to confer long term tumor regression due to a poor ability of CAR T cells to survive and infiltrate within a solid tumor mass. This may be partially explained by the tendency of CAR T cells to undergo functional exhaustion, similar to endogenous T cells (Chen *et al*., 2019). However, whether a PD-1-dependent downregulation of Drp1 activity is present also in exhausted CAR T cells is still not known. Should such modulation be observed, targeting Drp1 activity in CAR T cells (either pharmacologically or genetically) could represent a new strategy to ameliorate CAR T cell survival or infiltration.

To conclude, the modulation of Drp1 in tumor-derived exhausted T cells may represent a valuable target to ameliorate anti-cancer immune response in a number of instances, and the manipulation of CAR T system to this aim may represent a valid future strategy.

## 4. MATERIALS AND METHODS

### 4.1 Human samples

Peripheral blood samples were purified from buffy coats of healthy volunteer blood donors (independently of sex and age) under procedures approved by Institutional Review Board of Bambino Gesù Children’ Hospital (Approval of Ethical Committee N° 1314/2020 prot. N° 19826), including informed consensus for research purpose. Blood cells were incubated with RosetteSep Human T cell enrichment cocktail antibody mix (StemCell 15061). Unlabeled Human Peripheral Blood T (hPBT) cells were isolated by density gradient over Lymphoprep (StemCell 07811), with centrifugation for 20min at 1200rcf. Then T cells have been collected, washed and used for subsequent analyses. Human colon adenocarcinoma tissue sections were collected from the archives of the Tumor Immunology Laboratory, Department of Health Science according to the Helsinki declaration and under the approval of the University of Palermo Ethical Review Board (Approval N° 09/2018).

### 4.2 Mice

WT and Drp1^fl/fl^Lck:cre+ c57BL/6 mice were bred and maintained under conventional conditions at the Plaisant Srl (Castel Romano) Animal Facility. Drp1^fl/fl^Lck:cre+ mouse strain has been previously described (Simula *et al*., 2018). Mice were kept in cages of no more than 5-6 mice each, divided by sex, under 12h/12h light/dark cycles, with standard temperature, humidity and pressure conditions according to FELASA guidelines. Small red squared mice house and paper were used for cage enrichment. Mice health was monitored daily by veterinary staff and health analysis for pathogens were performed every three months according to FELASA guidelines. All mice were sacrificed by neck dislocation at 2-3 months of age. All efforts were made to minimize animal suffering and to reduce the number of mice used, in accordance with the European Communities Council Directive of 24 November 1986 (86/609/EEC). The mice protocol has been approved by the Allevamenti Plaisant Srl Ethical Committee as well as by the Italian Ministry of Health (Authorization #186/2020-PR). It has been written following the ARRIVE Guidelines, and the numeric details have been chosen following the criteria described in The National Centre for t he Replacement, Refinement and Reduction of Animals in Research (NC3Rs) (http://www.nc3rs.org.uk/). Sample size for the experiments performed has been established using power analysis method. Experiments involving growth of tumor cells in mice were performed using male mice.

### 4.3 Cell cultures and Reagents

Human Peripheral Blood T (hPBT) cells have been cultured in RPMI 1640 medium (Thermo Fisher 21875) supplemented with 10% Fetal Bovine Serum (Thermo Fisher 10270), 2mM L-glutamine (Thermo Fisher 25030081), 100U/ml penicillin/streptomycin (Thermo Fisher 15140130), 1x GIBCO MEM Non-essential amino-acids (Thermo Fisher 11140035), 1mM Sodium pyruvate (Thermo Fisher 11360039), and 100mg/ml Gentamycin (Thermo Fisher 15750045).

Murine T cells have been isolated from spleen using 70mm Cell Strainers (Corning 431751) and cultured in the same medium used for hPBT cells (complete RPMI medium) with the only exception of 50µM β-mercaptoethanol (Thermo Fisher 31350-010) addition.

MC38 tumor cells have been cultured in complete DMEM medium (Thermo Fisher 41966052) supplemented with 10% Fetal Bovine Serum (Thermo Fisher 10270), 2mM L-glutamine (Thermo Fisher 25030081), 100U/ml penicillin/streptomycin (Thermo Fisher 15140130), 1x GIBCO MEM Non-essential amino-acids (Thermo Fisher 11140035), 1mM Sodium pyruvate (Thermo Fisher 11360039), and 50µM β-mercaptoethanol (Thermo Fisher 31350-010).

Murine T cells have been isolated from spleen of WT mice and purified using Pan T Cell Isolation Kit (Miltenyi 130-095-130) or naïve CD8+ T Cell Isolation Kit (Miltenyi 130-096-543).

For *in vitro* activation, 2×10^5^ murine T cells have been stimulated with 5µg/ml anti-CD3 (plate-coated) (eBioscience 14-0031-86) and 1µg/ml anti-CD28 (Invitrogen 14-0281-86) for up to 48h in 96well plate. Alternatively, 5×10^5^ murine or human T cells have been stimulated at 1:1 ratio in 48well plate in presence of Sulfate Latex 4% w/v 5µm Beads (Molecular Probes S37227). For each experiment, 20×10^6^ beads were coated o.n. at 4°C with 2µg anti-CD3 (mouse: eBioscience 14-0031-86; human: eBioscience 16-0037-85) and 1µg anti-CD28 (mouse: Invitrogen 14-0281-86; human: 16-0289-85) and either 7µg of Recombinant PD-L1/B7-H1 Fc Chimera Protein (mouse: R&D System 1019-B7; human: R&D System 156-B7) (indicated as anti-CD3/28-PDL1-beads) or 7µg of Bovine Serum Albumin (Sigma A2153) (indicated as anti-CD3/28-beads). Cells defined as unstimulated were cultured in presence of beads coated with BSA only (o.n. coating of 20×10^6^ beads with 10µg of BSA). To modulate mTOR and ERK signaling, activated T cells have been incubated with 10nM RAD-001 (Novartis Oncology), 30µM FR180204 (Tocris 3706; indicated in the figures as ERKi) or 10 µM C6-Ceramide Cell-permeable ceramide analog (BML-SL110 Enzo Life Sciences). To inhibit autophagy, 20µM chloroquine (Sigma C6628) have been added to cells 1h before protein extraction.

To induce isolated murine naïve CD8+ T cells into an exhaustion-like state *in vitro*, cells have been stimulated up to 4 times with 5µg/ml anti-CD3 (plate-coated) (eBioscience 14-0031-86) and 1µg/ml anti-CD28 (Invitrogen 14-0281-86) for 24h in 96well plate. Between each stimulation, cells have been expanded using 20ng/ml mouse IL-2 (R&D System 402-ML). Cells were considered into exhaustion-like state after 4 stimulations (T_ex_) and were compared with effector-like (T_eff_) cells isolated from sibling mice and stimulated only once. For *in vitro* migration experiments, T_ex_ and T_eff_ cells were finally expanded *in vitro* for additional 6 days in IL2-containing medium and then used for the assays.

### 4.4 Western Blot

Western blot were performed as previously described (Simula *et al*., 2018). The following primary antibodies have been used: anti-actin (Cell Signaling 4970), anti-Drp1 (BD Bioscience 611113), anti-pS616-Drp1 (Cell Signaling 4494), anti-pS637-Drp1 (Cell Signaling 6319), anti-Mfn2 (Abcam ab56889), anti-Mfn1 (Santa Crux sc-50330), anti-Opa1 (BD Bioscience 612607), anti-Fis1 (Abcam ab71498), anti-Mff (Abcam ab129075), anti-pT202/204-ERK1/2 (Cell Signaling 4370), anti-ERK1/2 (Cell Signaling 4695), anti-Hsp90 (Cell Signaling 4877), anti-pSer2481-mTOR (Cell Signaling 2974), anti-mTOR (Cell Signaling 2983), anti-MnSOD (Enzo Life Sciences ADI-SOD-110), anti-LC3B (Cell Signaling 3868), anti-cFos (Cell Signaling 4384), and anti-GAPDH (Cell Signaling 2118). All primary antibody incubations were followed by incubation with appropriated secondary HRP-conjugated antibodies (GE Healthcare or Cell Signaling) in 5% milk plus 0.1% Tween20 (Sigma P2287). Detection of protein signals was performed using Clarity Western ECL substrate (Biorad 170-5061) and Amersham Imager 600. Stripping of the membranes for re-probing has been performed using buffer containing 1% Tween-20 (Sigma P2287), 0.1% SDS (Sigma 71729), and 0.2M glycine (VWR M103) at pH 2.2 (two washes for 10min).

### 4.5 Immunofluorescence

Immunofluorescence staining has been performed as previously described (Simula *et al*., 2020). Anti-TOM20 (Santa Cruz sc-11415) primary antibody was used to identify the mitochondrial network. Nunc Lab-Tek Chamber Slides (Thermo Fisher 154534) have been used to culture *in vitro* T cells directly on slides before fixation and were coated with 10ng/ml fibronectin (Millipore FC010) for 1h at RT before adding the cells. Images were acquired using a Perkin Elmer Ultraview VoX microscope. The mitochondrial network has been always evaluated upon 0.4mm slices z-stack reconstructions.

### 4.6 Immunohistochemistry

Formalin-fixed and paraffin embedded (FFPE) tissue samples of human colon cancer moderately differentiated (G2) cases were selected for in situ immunophenotypic analyses. Four-micrometres-thick sections were deparaffinized, rehydrated and unmasked using Novocastra Epitope Retrieval Solutions pH 9 in a thermostatic bath at 98°C for 30 minutes. Subsequently, the sections were brought to room temperature and washed in PBS. After neutralization of the endogenous peroxidase with 3% H2O2 and Fc blocking by a specific protein block (Leica Novocastra), the samples were incubated with phospho-DRP1 (Ser 616) (clone D9A1 Cell Signaling, 1:100) and PD-1 (clone NAT105 Abcam, 1:50) antibodies. IHC staining was revealed using MACH 2 Double Stain 1 kit (Biocare) and DAB (3,3’- Diaminobenzidine, Novocastra) and Vulcan Fast Red as substrate-chromogens. Triple immunostainings were performed by incubating the same sections with CD8 antibody (clone 4B11, 1:50 pH9, Leica Novocastra) and anti-mouse Alexa Fluor 488-coniugated secondary antibody (1:500, Life Technologies). Slides were analysed under a Zeiss Axioscope A1 and microphotographs were collected using a Zeiss Axiocam 503 Color with the Zen 2.0 Software (Zeiss).

### 4.7 Flow Cytometry

The following antibodies have been used to stain extracellular proteins: anti-CD8-Alexa488 (Biolegend 100723), anti-PD1-PE (eBioscience 12-9981-83), anti-CD4-PECy7 (Biolegend 100422), antiCD45-BV650 (Biolegend 103151), anti-CD44-BV421 (Biolegend 103039). Foxp3 Transcription Factor Staining Buffer Set (00-5523-00, eBioscience) has been used to stain intracellular proteins, detected with the following antibodies: anti-Drp1 (BD Bioscience 611113), anti-pS616-Drp1 (Cell Signaling 4494), anti-pS637-Drp1 (Cell Signaling 6319), anti-Mfn2 (Abcam ab56889), anti-Mfn1 (Santa Crux sc-50330), anti-Opa1 (BD Bioscience 612607), anti-Fis1 (Abcam ab71498), anti-Mff (Abcam ab129075), secondary goat anti-rabbit Alexa647 (Invitrogen A21244), secondary goat anti-mouse Alexa647 (Jackson 115-605-146), secondary goat anti-rabbit Alexa405 (Invitrogen A31556). Primary antibodies were incubated o.n. at 4°C, while secondary antibodies for 1h at RT. Background signals obtained by staining solely with secondary antibodies were subtracted from the corresponding signal from primary antibodies in the same cells to obtain MFI values reported in the figures.

To evaluate IFNγ production in tumor-derived T cells, cells have been restimulated for 4h in presence of 50ng/ml PMA (Sigma 79346), 1μg/ml ionomycin (Sigma I9657) and 2μM monensin (added for the last 2h, Sigma M5273) and then fixed and processed using Foxp3 Transcription Factor Staining Buffer Set (00-5523-00, eBioscience) and anti-IFNg-PE antibody (eBioscience 12-7311-82). Acquisitions have been performed using BD Accuri C6 and BD FACSCelesta cytometers. Cell sorting have been performed by staining cells with the aforementioned extracellular antibodies and using BD FACS Aria III flow cytometer.

For the evaluation of the mitochondrial membrane potential, 1μM TMRE (Thermo Fisher T669) has been added for 20 min and then the cells were washed and analysed. As a positive control for mitochondria depolarization, cells have been pre-treated with 50μM FCCP (Sigma Aldrich C2920).

### 4.8 Tumor induction

5*10^5^ MCA38 cells were injected subcutaneously into the right flank of two months-old male WT or Drp1^fl/fl^ and Drp1^fl/fl^Lck:cre+ mice. Mice were kept for up to 17/18 days in animal facility, and tumor growth was monitored twice or three times per week and recorded as [longest diameter]*[shortest diameter]^2^ in cubic millimetres. At days 7, 9, 11, 14 and 16 from tumor inoculation, mice were inoculated i.p. with 150µg of InVivoMab anti-mouse PD-1 (CD279), clone RMP1-14 (Bioxcell, BE0146) or 150µg InVivoMab rat IgG2a isotype control, clone 2A3 (Bioxcell, BE0089) antibodies (in 150µl of saline). Mice were randomly subdivided into each experimental group (a-IgG or a-PD1) before inoculation of antibodies (no specific randomization method was used). At day 17, mice were sacrificed and tumors were collected. Tumor tissues were mechanically dissociated over 70 mm-cell strainers, and mononuclear cells were enriched from tumor-derived cell suspensions by 40%/80% Percoll (GE Healthcare GE17-0891-01) density gradient, by collecting cells at the interface between 40% and 80% Percoll solution.

Isolated TILs have been used for subsequent proliferation and migration *in vitro* analyses or stained for flow cytometric measurements.

### 4.9 Seahorse analysis

Basal OCR has been measured in T cells during acute phase (unstimulated or stimulated for 12h) or after 48h of stimulation and 4 days of *in vitro* expansion with IL2, as previously described (Simula *et al*., 2018).

### 4.10 *In vitro* proliferation and migration assays

To evaluate the *in vitro* proliferation potential of tumor-infiltrated CD8+ T cells isolated from tumor-bearing Drp1^fl/fl^ and Drp1^fl/fl^Lck:cre+ mice, 2×10^5^ isolated TILs have been cultured *in vitro* in 96well plate in presence of 20ng/ml mouse IL-2 (R&D System 402-ML), 20ng/ml mouse IL-7 (R&D System 407-ML) and 20ng/ml mouse IL-15 (R&D System 447-ML). The total number of cells were estimated each 2/3 days using BD Accuri C6 flow cytometer and the percentage of CD8+ T cells in each plate evaluated by flow cytometry staining with anti-CD8-A488 (Biolegend 100723) antibody.

For transwell migration assays, 5×10^5^ TILs or *in vitro*-induced T_eff_ and T_ex_ have been starved from serum for 2h (by replacing FBS in the medium with 0.5% Bovine Serum Albumine, Sigma A2153) and then loaded on 5mm-pore size transwell filters (Costar 3421) and allowed to migrate for 2h in presence of 50nM CCL19 (R&D System 440-M3), 50nM CCL21 (R&D System 457-6C) or 10% Fetal Bovine Serum (Thermo Fisher 10270).

For the polarization assay, 2×10^5^ cells have been starved from serum for 2h (by replacing FBS in the medium with 0.5% Bovine Serum Albumine, Sigma A2153). Then, cells were allowed to adhere to 10mg/ml fibronectin-coated (Millipore FC010) microscope slides (Thermo Fisher ER302W-CE24) for 30min and stimulated by adding 50nM CCL19 (R&D System 440-M3) and 50nM CCL21 (R&D System 457-6C) for 15min before fixation and immunostaining. Alternatively, after serum starvation, 2×10^6^ cells were kept in 1.5ml Eppendorf tube and stimulated by adding 50nM CCL19 (R&D System 440-M3) and 50nM CCL21 (R&D System 457-6C) for 15min before directly proceeding to protein extraction.

### 4.11 Statistical Analysis

In the Figure legends, “n” indicates either the number of independent experiments (*in vitro* primary cells) or the number of mice used. Data are expressed as mean ± SEM from at least three independent experiments unless specified otherwise (Microsoft Office Excel and SigmaPlot v12.5 have been used for analysis). The number of mice used has been estimated using the power analysis method. All the acquisitions of the experiments have been performed blinded without knowing the specific conditions of each sample. Comparisons between groups were done using two-tailed Student’s T-test (two groups) or One-way and Two-way ANOVA (multiple groups and repeated measurements, adjustments for pairwise comparisons were performed using Holm-Sidak method). Mann-Whitney Rank Sum Test or ANOVA on ranks have been used if samples did not meet assumptions of normality and/or equal variance. Chi-square test has been used to evaluate data in Figure 1b. P-values are indicated in the Figures as follows: * = p < 0.05, ** = p < 0.01, *** = p < 0.001.

## 6. ACKNOLEDGEMENTS

This work was funded by Fondazione AIRC (Grant IG-2017 19826) to SC. LS was supported by a Fondazione AIRC “Fellowship for Italy” (23926).

## 7. AUTHORS’ CONTRIBUTION

LS and SC conceived the work. LS performed most of the experiments. YA helped with western blots and flow cytometry. VC and CT performed IHC on human tumor tissue. AC, CP, and GM performed seahorse analysis. SC raised funding. LS and SC wrote the manuscript, which has been approved by all authors.

