## Supplemental Material for "PD-1-induced T cell exhaustion is controlled by a Drp1-dependent mechanism"

a

b

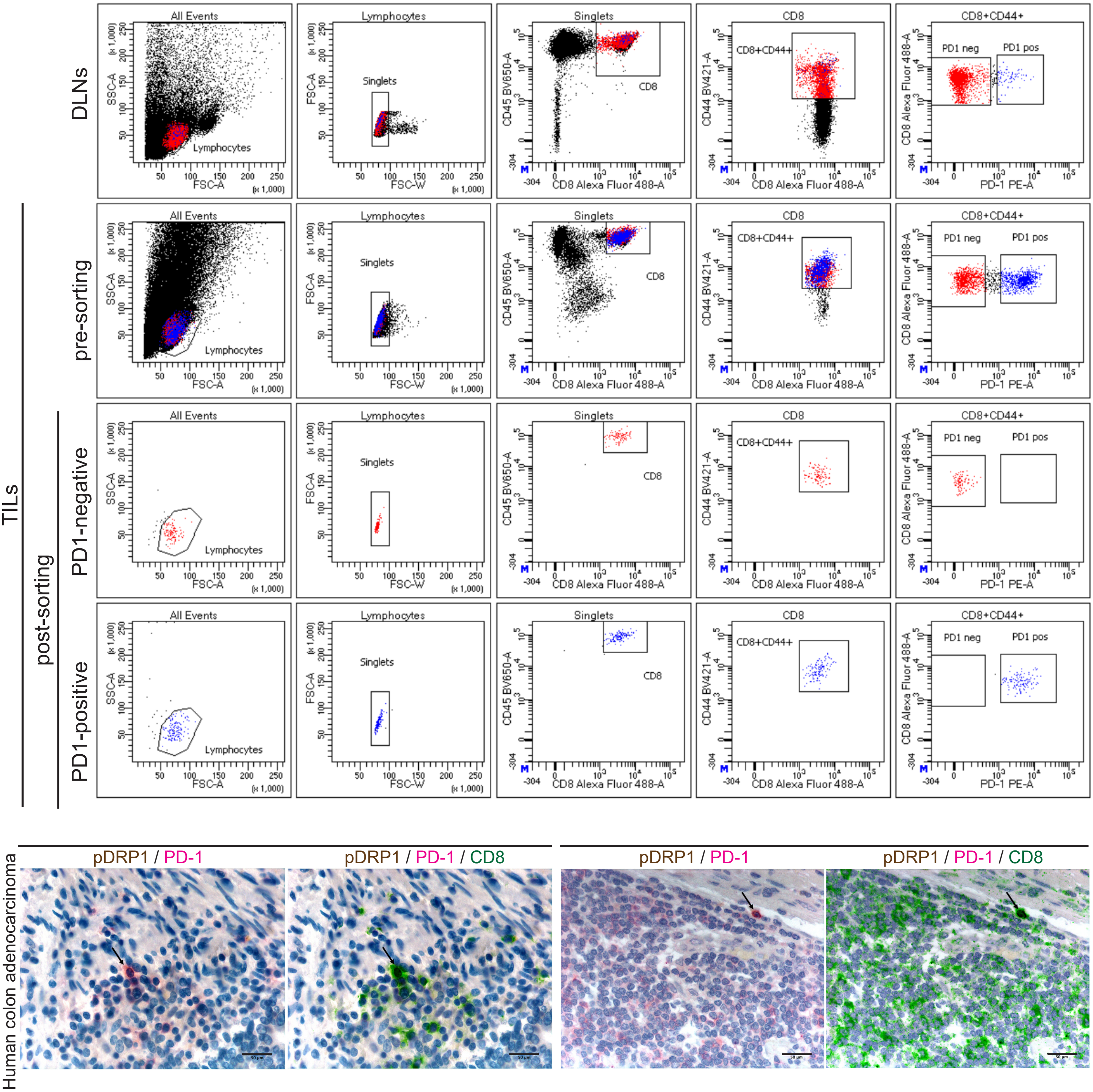


**Supplementary Figure 1. Gating strategy to isolate CD8⁺ TILs. Related to Figure 1.**

(**a**) Gating strategy to isolate CD45⁺CD8⁺CD44⁺ PD1$^{neg}$ and PD1$^{pos}$ T cells (used in Fig. 1b,c) from MC38-derived s.c. tumor masses grown for 18 days in WT c57BL/6 mice.

(**b**) Representative microphotographs of triple-marker immunohistochemistry for PD1 (rose), Drp1-pSer616 (pDrp1, brown) and CD8 (green) expression in lymphoid elements infiltrating human colon cancer.

Scale bar = 50µm.

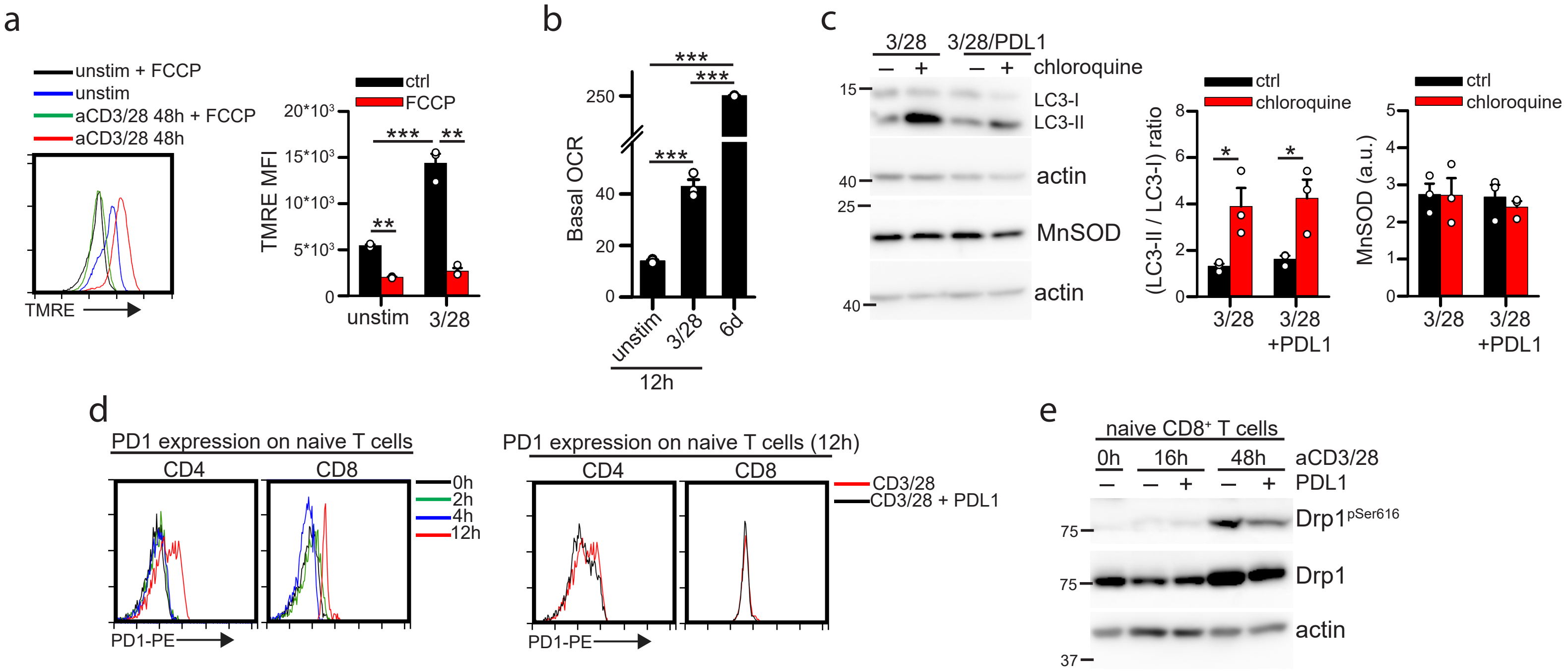


**Supplementary Figure 2. Functional analyses on activated T cells. Related to Figure 2.**
(**a**) TMRE profile of murine WT T cells left unstimulated or stimulated for 48h with aCD3/28-coated beads. FCCP has been used as a positive control for depolarized mitochondria. Quantification of the TMRE mean fluorescence intensity (MFI) is reported in the graph on the right (n=3).
(**b**) Basal oxygen consumption rate (OCR) measured by seahorse in WT murine T cells unstimulated (12h unstim) or stimulated for 12h (12h 3/28) or stimulated for 48h and then expanded for 4 days in IL2-containing medium (6d) (n=3).
(**c**) Murine WT T cells have been isolated and stimulated in presence of anti-CD3/28- or anti-CD3/28+PDL1-beads for 48h. Chloroquine has been added for the last hour in culture. The relative expression level of the indicated proteins is shown on the left. Quantifications of (LC3-II / LC3-I) ratio and MnSOD amount for each conditions are reported in the graphs on the right.
**(d)** Evaluation of PD1 expression by flow cytometry in purified murine CD4$^{+}$ and CD8$^{+}$ naive T cells stimulated *in vitro* with beads coated with anti-CD3/CD28 (left) or anti-CD3/CD28 with or without PDL1 (right) for the indicated time.
**(e)** Representative western blot images showing Drp1 and Drp1$^{pSer616}$ expression at the indicated time after stimulation in presence of anti-CD3/28- or anti-CD3/28-PDL1-beads in naive CD8+ T cells.
Data are shown as mean ± SEM. Significance is indicated as follows: * = p<0.05; ** = p<0.01; *** = p<0.001.

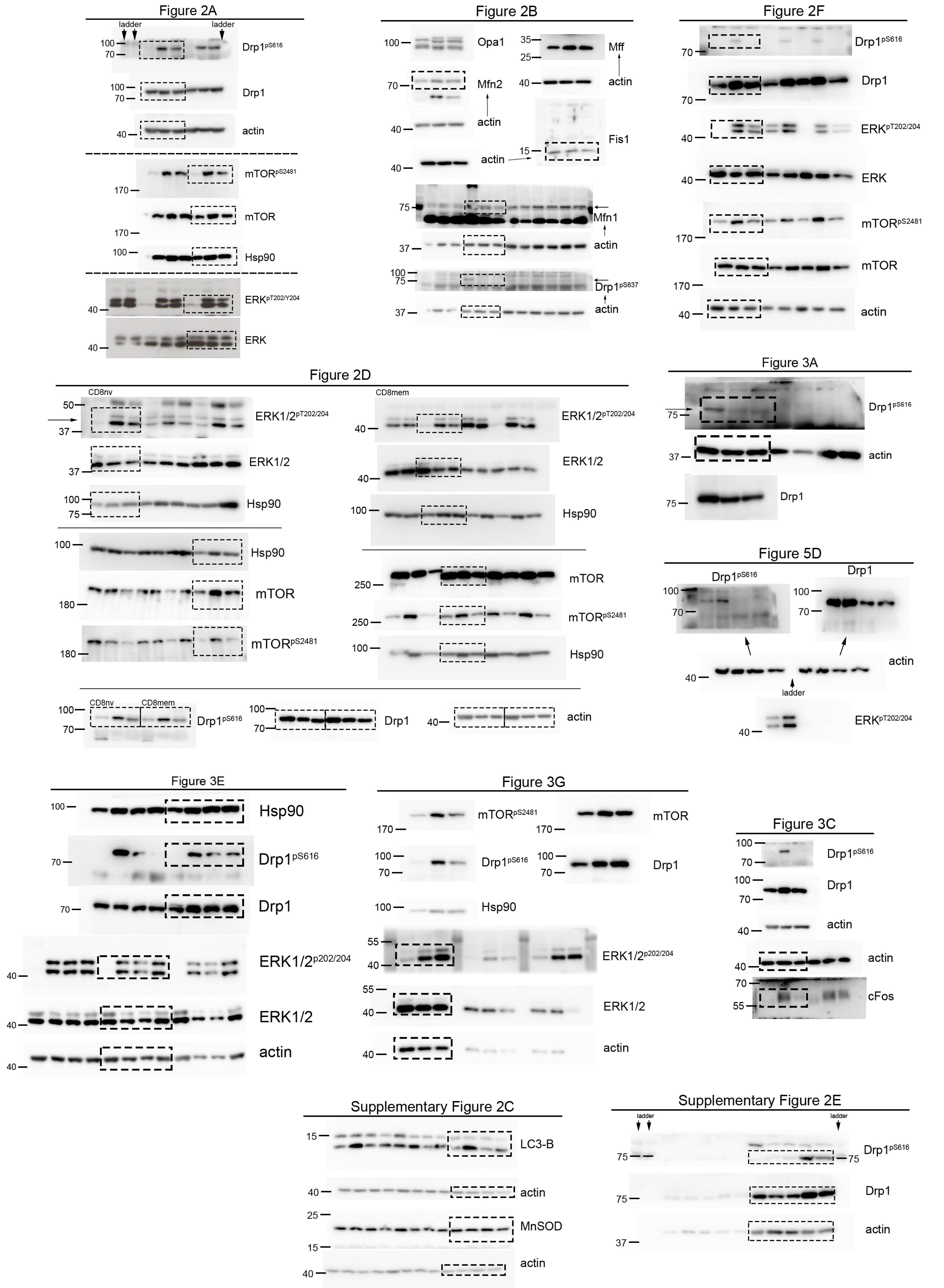


**Related Manuscript File.**

Uncropped images for all blots and gels in the manuscript.
